# Vargas: heuristic-free alignment for assessing linear and graph read aligners

**DOI:** 10.1101/2019.12.20.884676

**Authors:** Charlotte A. Darby, Ravi Gaddipati, Michael C. Schatz, Ben Langmead

## Abstract

Read alignment is central to many aspects of modern genomics. Most aligners use heuristics to accelerate processing, but these heuristics can fail to find the optimal alignments of reads. Alignment accuracy is typically measured through simulated reads; however, the simulated location may not be the (only) location with the optimal alignment score. Vargas implements a heuristic-free algorithm guaranteed to find the highest-scoring alignment for real sequencing reads to a linear or graph genome. With semiglobal and local alignment modes and affine gap and quality-scaled mismatch penalties, it can implement the scoring functions of commonly used aligners to calculate optimal alignments. While this is computationally intensive, Vargas uses multi-core parallelization and vectorized (SIMD) instructions to make it practical to optimally align large numbers of reads, achieving a maximum speed of 456 billion cell updates per second. We demonstrate how these “gold standard” Vargas alignments can be used to improve heuristic alignment accuracy by optimizing command-line parameters in Bowtie 2, BWA-MEM, and vg to align more reads correctly. Source code implemented in C++ and compiled binary releases are available at https://github.com/langmead-lab/vargas under the MIT license.

## 1 Introduction

Biological gold standards such as the Platinum Genomes (Eberle *et al*., 2017), synthetic diploid (Li *et al*., 2018), and Genome in a Bottle (Zook *et al*., 2014) catalog the variants present in a genome and are used to benchmark variant calling algorithms on real sequencing data. For benchmarking and algorithm development, using gold standard call sets is more realistic than simulating sequencing reads from a synthetic genome with known variants. However, read alignment algorithms, which determine a sequencing read’s point of origin with respect to a reference genome, are instead often evaluated using simulated sequencing reads due to the lack of a biological gold standard that directly answers questions about where sequencing reads should align.

In a typical read-aligner benchmark, sequencing errors and genetic variation are added to a substring of the reference genome, which is mapped back to the reference genome (Smolka *et al*., 2015). The simulated coordinate is then compared to the aligned coordinate. If the coordinates are equal (or within a predetermined range), the alignment is considered correct. We refer to this definition as “correct-by-location.” Critically, this procedure assumes that the desired (optimal) alignment is at the same location where the read was simulated, which may not be the case due to the simulation process and genomic repetitiveness. Furthermore, the base-level alignment may be suboptimal even if the read is mapped to the simulated location; e.g. the alignments may disagree about the exact placement and number of gaps in the alignment. Additionally, simulated reads may not faithfully capture more subtle aspects of the sequencing processing, especially any sequence-specific error modalities (Aird *et al*., 2011).

A central component of all read aligners is defining a scoring function. The scoring function penalizes differences between the read and reference; i.e. it is the objective function that the aligner seeks to optimize. When the optimal alignment score for a read equals the score reported by the algorithm, we call this alignment “correct-by-score.” Optimal alignments of real sequencing reads could be considered a computational gold standard for read alignment algorithms (c.f. Holtgrewe et al. (2011)). These can be used to evaluate alignment algorithms using real data in the same way as biological gold standard variant catalogs are used to assess variant callers. To serve as a computational gold standard, the optimal alignment should be calculated with respect to the same alignment mode (local or semiglobal), scoring function, and reference genome of the algorithm being evaluated.

While most current heuristic and heuristic-free read alignment algorithms assume that the reference genome is linear, with greater understanding of genetic diversity has come increasing focus on alternatives to the linear reference genome. Various solutions have been proposed that incorporate information about genetic variation in the population, including graph-shaped reference genomes (Paten *et al*., 2017), pan-genomes (Yang *et al*., 2019), and a genome that contains the most common (major) allele at each variable site (Pritt *et al*., 2018; Ballouz *et al*., 2019). The most recent human reference genome assembly, GRCh38, includes alternate assemblies for hypervariable loci (Church *et al*., 2015). Alignment algorithms that account for genetic variants have likewise been proposed (Schneeberger *et al*., 2009; Vijaya Satya *et al*., 2012; Huang *et al*., 2013; Garrison *et al*., 2018; Rautiainen *et al*., 2019; Jain *et al*., 2019).

Whether the genome is linear or a variant graph, calculating the optimal read alignment using dynamic programming is extremely work-intensive. Popular algorithms for aligning to linear genomes such as Bowtie 2 (Langmead and Salzberg, 2012) and BWA-MEM (Li, 2013), or to graph genomes, such as HISAT2 (Kim *et al*., 2019) and vg (Garrison *et al*., 2018), scale to genomic-scale datasets because they use heuristics to find approximate solutions to the optimization problem defined by the objective function. A widely used heuristic is “seed-and-extend” in which short exact sequence matches are used to anchor the search for longer matches. Relatedly, aligners apply various heuristics to prioritize which candidate seed matches are most likely to lead to a high quality alignment. These computational shortcuts limit effort spent on candidate alignments that appear like they will be suboptimal, but may cause the algorithm to return a suboptimal alignment or no alignment.

Alignment errors due to heuristics can ultimately lead to confounding of scientific results downstream. At the same time, it is difficult to study the precise effects of heuristics. This is both because they are varied and complex, and because it is computationally demanding to disentangle the effects of heuristics from those of scoring functions and other read-alignment parameters. A case in point is the Qtip study (Langmead, 2017), where mapping-quality predictions were improved by modifying the source code of heuristic alignment algorithms to report specific information about the heuristic search procedure for each read. Because a computational gold standard can be constructed to match the aligner’s scoring function and other parameters, it uniquely allows us to isolate the effects of heuristics.

General-purpose computer processors have recently become capable of running hundreds of separate threads of execution simultaneously in parallel. Within each thread, single-instruction multiple data (SIMD) (“vector”) operations can be employed that operate on a group of values at once instead of a typical processor instruction that operates on individual scalar values. Vargas implements a form of the Smith-Waterman (Smith and Waterman, 1981) algorithm for query-reference sequence alignment to a linear reference or a directed acyclic graph (DAG). Rognes (2011) proposed a query-parallel strategy that fills the same element (same row and column) in several dynamic programming matrices at once. This has been shown to be efficient and practical in prior studies (Rahn *et al*., 2018; Jain *et al*., 2019). We adopt this approach in Vargas and provide further evidence that it is efficient on modern general-purpose architectures for both linear and graph alignment.

Vargas is the most efficient and flexible tool for establishing computational gold standards for evaluating read alignment heuristics and scoring schemes. While the need for such standards was discussed and partially addressed in the previous Rabema study (Holtgrewe *et al*., 2011), Vargas is an advance over those methods owing to its handling of graph-shaped as well as linear reference genomes, efficient scaling to many threads, use of modern SIMD instructions, and handling of flexible affine-gap-penalty scoring schemes such as those used in modern aligners like vg, Bowtie 2 and BWA-MEM. Using RNA-seq reads aligned to the transcriptome, we can also assess the accuracy of alignment scores reported by the pseudoaligner Salmon. We demonstrate the utility of the method by showing how it can be used to assess alignment correctness and mapping quality predictions, and to optimize Bowtie 2, BWA-MEM, and vg alignment parameters for whole-genome sequencing and ChIP-seq reads. Importantly, in these cases the analyses are guided by real reads.

## 2 Materials and methods

Vargas is a heuristic-free read aligner for linear and graph reference genomes. Among currently available software and libraries, only Vargas has the features required to calculate optimal alignments for semiglobal and local alignment to linear and graph genomes, with affine gap penalties and base-quality-dependent mismatch penalties. These optimal alignments of real sequencing reads, calculated with respect to the particular optimization problem each heuristic algorithm seeks to approximate, can be used to systematically evaluate the behavior of the heuristic tools and improve the algorithms. We provide the first thorough evaluation comparing graph alignment to linear-genome alignment without the confounding factors that come when comparing tools with differing scoring functions or heuristics and using real rather than simulated sequencing reads.

### 2.1 Graph Alignment

The Smith-Waterman algorithm uses dynamic programming to find the optimal local alignment of a query sequence to a reference sequence under a scoring function rewarding matches and penalizing mismatches and gaps (Smith and Waterman, 1981). Affine gap penalties (Gotoh, 1982) are often employed, where the penalty to start a gap is greater than the penalty to extend the gap, encouraging larger consecutive gaps rather than multiple smaller gaps. The algorithm was further extended to support local alignment of a linear query to a directed acyclic graph (DAG) reference (Lee *et al*., 2002). Genetic variants induce “forks” and “joins” in the DAG (Figure 1). The classical dynamic programming recurrence is computed for each column of each vertex in a topological-sort order, taking into account cases where a column has more than one predecessor in the graph structure (e.g. column 7 in Figure 1).

**Figure 1:**
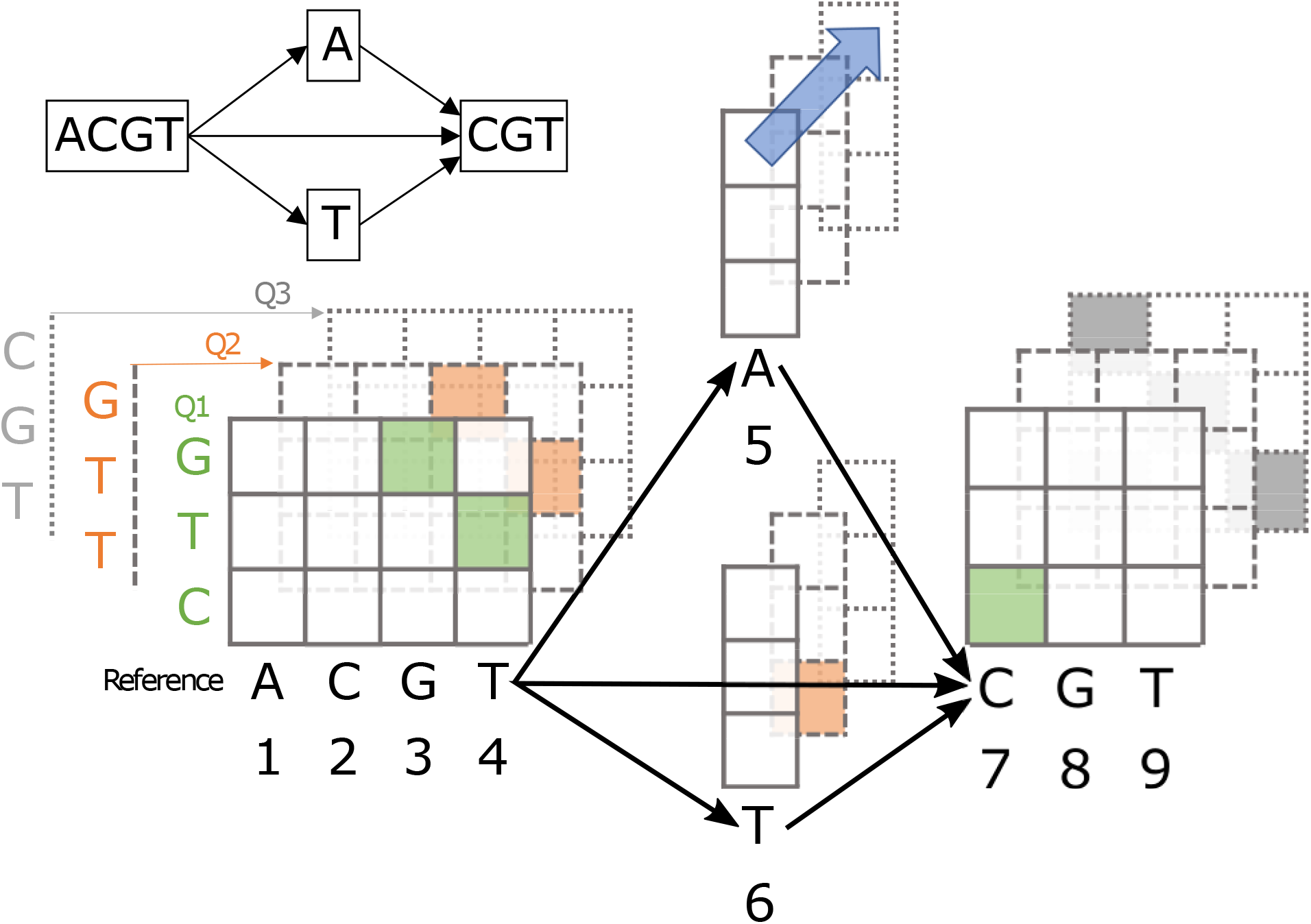
The multiple alignment of three sequences (ACGTACGT, ACGTTCGT, ACGTCGT) can be represented as a four-node graph. Suppose we want to compute the dynamic programming alignment matrix for three length-3 queries to the graph-shaped reference. The columns of the matrices correspond to characters in the reference and are numbered 1 through 9 in the order they will be computed in the dynamic programming algorithm. Query characters label the rows of each matrix. Matrices are shown stacked because SIMD instructions operate on the same row and column in multiple matrices simultaneously (e.g. the cells connected by the blue arrow). The optimal score for query Q1 is in column 7 (shaded cells show the alignment traceback); for Q2 the optimal score is in column 6; for Q3 two equally good alignments end in columns 4 and 9 (possible traceback is shown).

Before alignment, Vargas constructs the DAG from a linear reference sequence and optionally a set of genetic variants in a Variant Call Format (VCF) file. The graph can be computed once and is stored on disk in a format that includes the sequence represented by each node and links between nodes. To reconcile coordinate shifts introduced by insertions and deletions, alignments are anchored to the reference sequence. Nodes representing parallel paths with different sequence lengths are right-aligned to the reference sequence. Vargas produces read alignments in SAM format. For the best and second-best alignment scores, Vargas reports the reference position of the rightmost aligned query base and the count of equally-scoring alignment locations at least one read-length apart in custom SAM tags. For linear genomes, the alignment traceback can optionally be computed to populate the CIGAR and POS fields.

### 2.2 Vectorization Strategies

Vectorized dynamic-programming alignment strategies involve filling several cells of the dynamic-programming matrix at the same time. An early approach computed chunks of cells along the minor diagonal (Wozniak, 1997), a later approach achieved a more predictable memory-access pattern by filling vertical chunks (Rognes and Seeberg, 2000), and a “striped” variation on this idea achieved better performance by simplifying the loops (Farrar, 2007). In these later algorithms, elements in a chunk are inter-dependent because the final value of an element depends on the final values of neighbors above. Thus, the initial filling step must be followed by a “repair” step to resolve dependencies. As architectures evolve to allow more elements to be computed at once, the amount of work required for repair increases. These methods are therefore not well positioned to benefit from increases in vector width, such as those afforded by AVX512BW (byte and word) extensions that allow for simultaneous calculation of 64 8-bit matrix elements (Intel Corporation, 2015). Instead, Vargas uses the query-parallel strategy of Rognes (2011). In this paradigm, the value of each vector element is independent so no repair step is required. For a particular row and column position in the matrix, SIMD instructions are used to compute the value for 8–64 queries simultaneously, depending on the vector capacity. The query-parallel strategy has been used for alignment to linear references in SeqAn (Rahn *et al*., 2018) and to directed acyclic graphs in PaSGAL (Jain *et al*., 2019).

### 2.3 Implementation

Vargas is implemented in C++ using SIMD instructions and supports the SSE4.1, AVX2, and AVX512BW instruction sets, which can be compiled for many architectures including Intel Xeon Phi (Knights Landing/KNL) and Xeon Platinum (Skylake/SKX). The KNL architecture supports 256–288 threads across 64–72 cores (Sodani, 2015; Jeffers *et al*., 2016) and SKX can be configured with up to 28 cores, each with two AVX (advanced vector extensions) processors (Tam *et al*., 2018). In Vargas, to maximize throughput, each SIMD word (vector) is split into 8-bit operands allowing for the simultaneous alignment of 16, 32, and 64 sequences with 128-bit, 256-bit, and 512-bit vectors for SSE4.1, AVX2, and AVX512BW respectively. If the difference between the maximum and minimum possible alignment scores exceeds 255 based on the read length and scoring function, 16-bit operands are selected at runtime.

## 3 Results

First we used Vargas to study the performance and accuracy of heuristic read aligners on real sequencing data. While most benchmarks use simulated reads, using a heuristic-free aligner allows us to determine whether the alignment for a real read is correct-by-score, as others have observed (Holtgrewe *et al*., 2011). We explored how alignment settings can affect which reads are incorrect-by-location, incorrect-by-score, or completely fail to align due to heuristics. We evaluated the time-accuracy tradeoff of the Bowtie 2 and HISAT2 effort presets and propose a comparable set of parameters for BWA-MEM and BWA aln. Based on the correct-by-location definition, we compare BWA-MEM and Bowtie 2 mapping quality to the mathematical ideal before and after adjustment with Qtip (Langmead, 2017). Finally, we show how a small set of reads annotated with optimal alignment score using Vargas can be used to optimize Bowtie 2 alignment parameters and suggest a possible algorithmic improvement. Data and scripts to reproduce the experiments in this section and the next are available at https://github.com/cdarby/vargas-experiments.

### 3.1 Computational performance

We measured performance using GCUPS (giga cell updates per second), which is the number of cells in the dynamic programming matrix computed per second. This standard metric used extensively in the literature normalizes performance to enable comparisons between experiments conducted using reads and references of different lengths (Rahn *et al*., 2018; Daily, 2016; Jain *et al*., 2019; Liu and Schmidt, 2014; Liu *et al*., 2013). Evaluation was performed on an Intel Xeon Phi 7250 (Knights Landing / KNL) computer with 68 cores and four threads per core, and an Intel Xeon Platinum 8160 (Skylake / SKX) computer with two 24-core processors and two threads per core. GCUPS results for semiglobal alignment are shown in Figure 2; GCUPS results for local alignment and wall time results are in Figures S1-S4. Ideal scaling would be linear. Instead we see less-than-ideal (sublinear) scaling owing to hyperthreading and, more generally, to contention for shared resources. SKX employs two-way hyperthreading on 48 physical cores, and speedup is observed up to 64 threads with AVX512BW instructions (Figure 2a-b). KNL employs four-way hyperthreading on each of its 68 physical cores, allowing a maximum of 272 simultaneous threads. With AVX2 instructions, the GCUPS performance doubles from 68 threads (one per core) to 136 threads (two per core) but does not continue to double when there are three or four threads per core (Figure 2c-d). See Supplementary Figures S1 and S2 for scaling results with local alignment.

**Figure 2:**
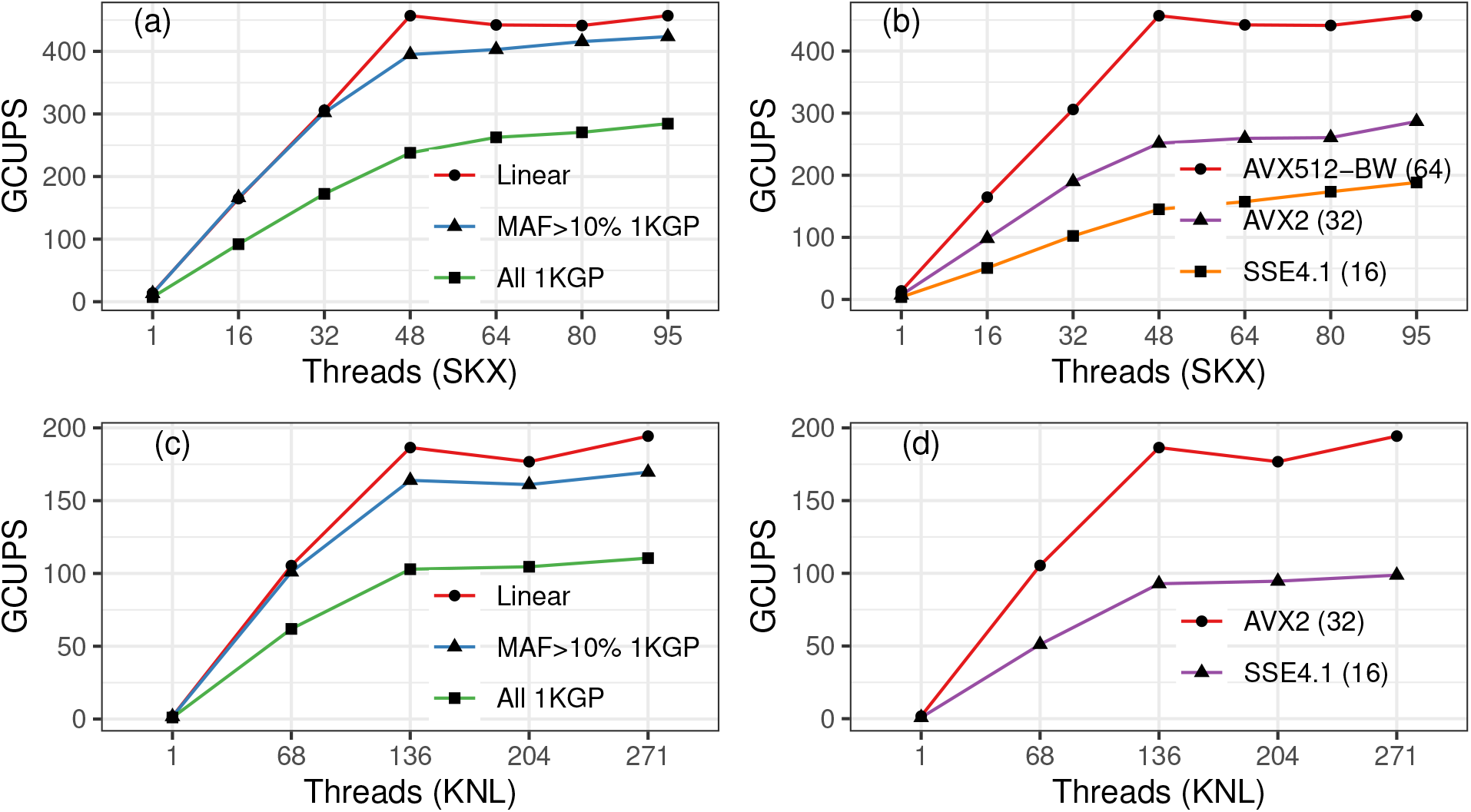
Scaling results for Skylake (SKX; a and b) and Knight’s Landing (KNL; c and d) architectures and the instruction sets available for each. The input number of reads is scaled in proportion to the number of threads, so that each thread aligns a full vector of 100bp reads against the reference 8 times. (a) and (c). Reads were aligned to chromosome 19 with no variants (Linear, 1 node), 1000 Genomes Project Phase 3 variants with minor allele frequency > 10% (MAF >10% 1KGP, 436K nodes), or all 2504 individuals’ variants from 1000 Genomes Project Phase 3 (All 1KGP, 5.1M nodes). (b) and (d). Reads were aligned to chromosome 19 with no variants. Vector size is shown in parentheses after the instruction set name on the right panels. Note, KNL does not support the AVX512BW instruction set. GCUPS = giga cell updates per second.

The best speed we observed using Vargas is 456 GCUPS, which was observed for semiglobal alignment to the linear reference of chromosome 19 using AVX512BW instructions with 64-way vectorization on 48 threads of the SKX computer. When the chromosome 19 graph contained all 1000 Genomes Phase 3 SNPs and indels (5.1 million nodes and 3.66% increase in linearized genome size) we observed 237 GCUPS with the same configuration. When aligning 150bp reads to chromosome 10, SeqAn (Rahn *et al*., 2018) reported 420 GCUPS using 40 threads on SKX with AVX512BW; on the same dataset, Parasail (Daily, 2016) recorded 74 GCUPS, but used AVX2 instructions that offer only half the throughput per instruction. PaSGAL (Jain *et al*., 2019), reported 317 GCUPS with 48 threads on SKX with AVX512 instructions when aligning 100bp reads to a graph genome of the 1Mbp Leukocyte Receptor Complex locus including all variants from 1000 Genomes Project Phase 3, not including traceback.

Vargas semiglobal alignment to the linear genome on Xeon Phi (KNL) with AVX2 instructions and 271 threads achieved a maximum speed of 194 GCUPS. Alignment to the all-variant graph described above achieved 110 GCUPS with the same configuration. This compares favorably to previous Xeon Phi-based efforts such as SWAPHI (58.8 GCUPS) (Liu and Schmidt, 2014), as well as to GPU based aligners such as CUDASW++3.0 (119 GCUPS, GTX680) (Liu *et al*., 2013). While several exact dynamic programming pairwise alignment algorithms are available in the literature (summarized in Supplementary Table S3), Vargas offers the most flexibility in terms of scoring function and options for local, semiglobal, linear, and DAG alignment with comparable speed and scaling to the state of the art.

### 3.2 Alignment Accuracy

We use 100,000 unpaired 100bp reads from the 1000 Genomes Project sample NA18505, SRA accession ERR239486, and 100,000 unpaired 250bp reads from the 1000 Genomes Project sample NA19017, SRA accession SRR1295544. All aligners were evaluated with respect to the human reference genome GRCh38 primary assembly. Since HISAT2 and vg have graph alignment capabilities, we also built graph-genome indexes for both using the 6.2 million SNPs and indels from the 1000 Genomes Phase 3 call set with allele frequency at least 10% (Lowy-Gallego *et al*., 2019). Vargas alignments to the linear genome took about 40 hours and alignments to the graph genome took about 57 hours, using 96 threads on a SKX computer with AVX512BW instructions and 64-way vectorization.

Figure 3 shows the performance of Bowtie 2 (Langmead and Salzberg, 2012), BWA-MEM (Li, 2013), HISAT2 with graph genome (Kim *et al*., 2019) and vg with graph genome (Garrison *et al*., 2018) on the 100bp read set with respect to the correct-by-score definition. Bowtie 2 and HISAT2 were run multiple times using their “preset” parameters that trade between runtime and alignment accuracy. We also determined a sequence of settings that create a similar tradeoff for BWA-MEM. Additional results for BWA aln (Li and Durbin, 2010), Bowtie 2 with local alignment (default is semiglobal), vg with linear genome, and HISAT2 with linear genome are in Supplementary Figure S3; plots evaluating alignments based on the correct-by-location definition are in Supplementary Figure S4.

**Figure 3:**
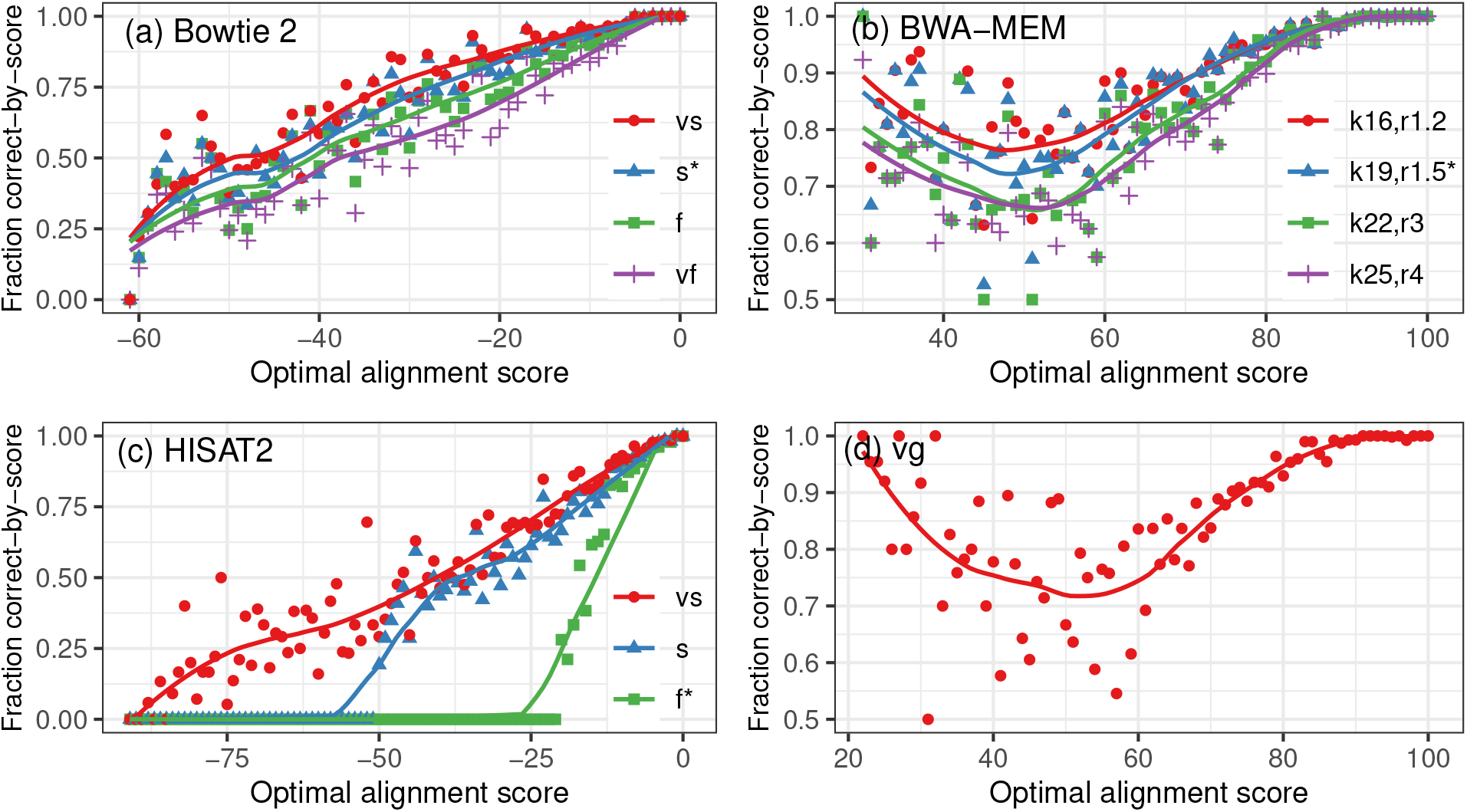
The 100,000 100bp read set is binned by the optimal alignment score calculated by Vargas, shown on the horizontal axis, which is truncated at the point after which no alignments are reported by the heuristic. A line is fitted to the scatterplot of fraction of reads that are correct-by-score. Only primary alignments were evaluated for HISAT2 and BWA-MEM. Default parameter settings denoted by star (*).

*Bowtie 2*. Bowtie 2’s default semiglobal alignment mode offers four “presets” providing four sets of values for the `-D` (extension effort) `-R` (re-seeding) `-L` (seed length) and `-i` (seed spacing) parameters. We found that these presets effectively trade between time and accuracy when evaluated using correct-by-score. For the settings that are faster but less accurate, the number of reads failing to align increases and the number of reads aligned correctly decreases. Correctness decreases approximately linearly with optimal alignment score (Figure 3a). Interestingly, when the optimal alignment score is below −40, approximately 50% of the reads are incorrect-by-score, meaning no alignment was reported or the heuristics have lead to an erroneous suboptimal alignment for the majority of these reads.

*HISAT2*. The latest version of HISAT2 (version 2.2.0-beta https://github.com/DaehwanKimLab/hisat2/tree/hisat2_v2.2.0_beta) provides two modes that change the `--score-min` (minimum score for reporting alignments) and `--bowtie2-dp` parameters to align more reads. `very-sensitive` uses “unconditional dynamic programming” and `sensitive` uses “conditional dynamic programming.” Compared to the default ‘fast’ mode, more reads are correct-by-score using the very-sensitive and sensitive modes, but more reads are incorrect-by-score. This is possible because more reads are aligned. In contrast, the slowest Bowtie 2 and BWA-MEM presets have the least incorrect reads (Supplementary Table S1). More incorrect alignments are likely reported because the HISAT2 presets also alter the minimum score of alignments reported, which is an independent parameter in Bowtie 2/BWA-MEM.

*BWA-MEM*. While BWA-MEM does not offer presets like Bowtie 2 and HISAT2, we selected four settings for `-k` (seed length) and `-r` (re-seeding) that trade between speed and accuracy. With the default parameter settings (blue line) and our proposed presets, correctness decreases with optimal score for reads with score greater than 50, but then increases with optimal score for reads with optimal score between 50 and 30, the minimum alignment score reported (Figure 3b). This appears to be because most of the aligned reads with optimal score below 50 have an exact or near-exact alignment that does not include all the bases in the read, sometimes referred to as “soft clipping.” The BWA-MEM heuristic algorithm for local alignment is often able to correctly identify optimal alignments of this sort, even though they have low alignment score. In contrast, illustrating a fundamental difference between the local and semiglobal alignment problems, the Bowtie 2 and HISAT2 strategies for semiglobal alignment seldom identify the optimal low-scoring alignments which have many mismatches and gaps because every base of the read must be included in the alignment.

*vg*. Like BWA-MEM, vg performs local alignment. Correctness decreases with optimal score for reads with score greater than 50 but increases with optimal score for reads with optimal score between 50 and 20, which again appears to be due to low-scoring local alignments frequently being soft-clipped to involve a short region of high similarity and underscores the differences between local and semiglobal alignment.

*Salmon*. The Salmon pseudoaligner (Patro *et al*., 2017) version 1.1.0 will optionally perform semiglobal dynamic programming alignment and produce a SAM file with alignment scores based on an edit distance scoring function with the command-line flags `--writeMappings --validateMappings`. While the alignments themselves are not designed to be used in downstream analysis (e.g. the CIGAR string does not correspond to the reported alignment score), the alignment scores are used in this mode to select which alignments will be used in expression quantification. Using an RNA-seq dataset and Vargas alignments to the reference transcriptome, described fully in the Supplementary Information, we evaluated the alignment scores reported by Salmon with the default algorithm which performs alignment only between and beyond the maximal exact matches (MEMs), and with the `--fullLengthAlignment` flag, which performs alignment along the whole read. The default algorithm had more correct-by-score alignments compared to the full-length algorithm (Supplementary Table S5, Supplementary Figure S6).

Supplementary Figure S4 compares the correct-by-score and correct-by-location definitions for the genomic alignments of 100bp real sequencing reads with unique and repetitive optimal alignments. When we consider only the reads that align uniquely, there is little difference between the definitions of correctness. When we consider reads that align repetitively, noticeably more reads are correct-by-score than are correct-by-location, as expected. We also performed Vargas alignments of 100,000 250bp reads for the same aligners (Supplementary Table S2, Supplementary Figure S5) and the results closely match those obtained with 100bp reads.

To compare the simulation-based definition of correct-by-location, which is calculated by comparing the genomic coordinate where the read is simulated from to the coordinate where it is aligned, and the correct-by-score definition, which is calculated using Vargas matched to the scoring function of the alignment algorithm, we generated simulated reads from human chromosome 19. For alignments with Bowtie 2 to the chromosome 19 reference, we saw that 7.79%, 2.80%, and 1.31% of reads of length 50, 100, and 150 respectively, were correct-by-score but incorrect-by-location. BWA-MEM produces similar results. Details of this experiment are in the Supplementary Information and Supplementary Table S4.

### 3.3 Mapping Quality

The mapping quality (MAPQ) of a read alignment is defined (Li *et al*., 2008) as:

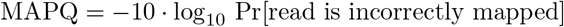

An accurate prediction for MAPQ requires an accurate prediction for the probability the alignment is incorrect. Heuristics make this difficult by effectively “censoring” the space of alignments the aligner can find. Because of this, heuristic aligners work with only partial information when making a MAPQ prediction. This leads to errors, such as predicting a high MAPQ for an incorrect alignment or for an alignment that truly deserves a low one. Since downstream tools such as variant callers depend on MAPQs to make decisions about how to weigh and filter evidence, it is important to predict accurately.

Aligners predict MAPQ based on features such as the alignment score of the best and second-best alignments found or how repetitive the seed hits are. It may also depend on the number of hits with the same score as the reported alignment (Li *et al*., 2008). To condense such features into a single score, BWA-MEM and vg use a formula, whereas Bowtie 2 and HISAT2 use a decision tree-like approach. Some aligners do not attempt to estimate mapping quality at all.

Qtip (Langmead, 2017) adjusts mapping quality using tandem simulation and extra output from the heuristic during the alignment algorithm. Using Vargas, we can assess how well mapping quality reflects alignment correctness by grouping reads by their aligner-assigned or Qtip-adjusted mapping quality and calculating average correctness. In this case, it is important to use the correct-by-location definition (within a 5 bp buffer), to match the definition of MAPQ. Figure 4 shows results for Bowtie 2 with semiglobal or local alignment and BWA-MEM (red lines), and Qtip-adjusted (blue line) along with a black line indicating where the points would lie if they conformed perfectly to the mathematical definition of MAPQ, for the 100bp and 250bp read sets. Consistent with past experiments, Qtip-adjusted mapping qualities fall into a smaller numerical range than aligner-calculated MAPQ and are generally more monotonic and closer to the ideal.

**Figure 4:**
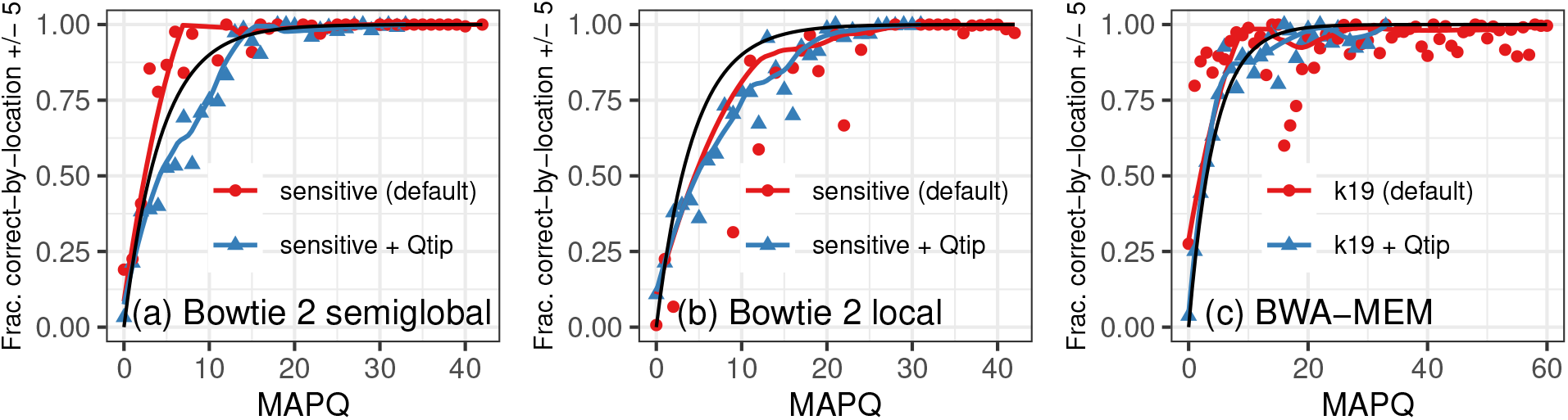
The 100bp read set (a-c) and 250bp read set (d-f) are binned by mapping quality, shown on the horizontal axis, and correctness is measured using the correct-by-location definition within 5bp. The black line reflects the mathematical definition of mapping quality.

**Table 1:** Bowtie 2 semiglobal alignment (SG), Bowtie 2 local alignment (L), BWA-MEM, and vg using MAF > 10% graph genome, were run on 8,365 difficult reads from the 100bp dataset with seed length varying from 10 to 35. The value in the table equals the ratio between the measurement (row) when using the stated parameter setting (column) versus when using default parameters. For BWA-MEM and vg, the second parameter setting is the one that gave the lowest mean AS difference without taking more than 1.5 times as long as the default.

|  | **Bowtie2 SG** | **Bowtie2 L** | **BWA-MEM** |  | **vg** |  |
| --- | --- | --- | --- | --- | --- | --- |
| Seed length | 28 | 32 | 14 | 17 | 10 | 12 |
| % aligned | 1.00 | 0.95 | 1.00 | 1.00 | 1.02 | 1.02 |
| % correct score | 1.01 | 1.02 | 1.00 | 1.00 | 1.01 | 1.01 |
| Mean AS difference (all aligned reads) | 0.82 | 0.90 | 0.93 | 0.98 | 0.61 | 0.66 |
| Mean AS difference<br>(incorrect-by-score aligned reads) | 0.89 | 1.02 | 0.97 | 1.00 | 0.81 | 0.85 |
| Alignment time | 0.76 | 0.56 | 1.97 | 1.20 | 1.99 | 1.41 |

## 4 Optimizing read alignment

As an example of how Vargas alignments can be used to improve the heuristic alignment workflow by a user who is not necessarily a tool developer, we further examined the 100bp reads described in Section 3.2, specifically the 8,365 reads that were unaligned by Bowtie 2 or had an alignment with >1 mismatch or at least 1 gap, with default parameters. All further analysis was performed on this subset of the original dataset, which we refer to as ‘difficult reads’.

We wanted to determine whether more accurate alignments of difficult reads could be obtained by tuning command-line parameters without much increase in runtime, so we varied the seed length parameter of each aligner from 10 to 32 (Supplementary Excel File 1). In Table 4 we report results for the default seed length and the seed length that minimized the average difference between the optimal alignment score (first column for each aligner) and the heuristic alignment score, for aligned reads (second column). Notably, the optimal seed length for Bowtie 2 in both semiglobal (SG) and local (L) alignment modes was faster than the default seed length. The optimal seed length for BWA-MEM and vg was slower than the optimal seed length, so we also included results for the optimal parameter that had runtime less than 1.5 times slower than the default parameters.

This case study demonstrates how a set of real reads annotated with the optimal alignment score can be used to tune heuristic alignment parameters that are exposed to the user on the command line. While a 1-2% increase in correctly-scored alignments may seem marginal, this would have a significant impact on a dataset with millions or billions of reads and on use cases with a low signal-to-noise ratio such as cell-free DNA analysis or somatic variant calling. The parameters enabling this increase in accuracy can be identified with Vargas alignments of just a few thousand reads.

We also performed a similar optimization experiment for Bowtie 2, BWA-MEM and vg with graph genome using ChIP-seq reads, described in the Supplementary Information. By varying command-line parameters on a small test set of 10,000 reads, we observed increased alignment rate and correctness-by-score on 570,000 difficult reads from the dataset, at the cost of increased runtime (Supplementary Table S6 and Supplementary Excel File 2).

## 5 Discussion

We presented Vargas, a heuristic-free read alignment tool achieving extremely high multi-threaded throughput. Vargas works with flexible alignment scoring functions (e.g. affine gap penalty) and parameters (e.g. local and semi-global alignment), and with both linear and graph references. Read alignments produced by Vargas can be used as a computational gold standard for evaluating short-read alignment algorithms, including with real sequencing datasets, and in much the same way as biological gold standards are used to assess variant calling algorithms. Vargas opens the door to comprehensive study of the effects of alignment heuristics and, distinct from that, the effects of alignment scoring functions. Though the default scoring functions of tools like BWA-MEM and Bowtie 2 are widely used, they are not very well studied, and this is in large part because it is difficult to separate the effect of the scoring function from the closely related effects of the heuristics. Vargas alignments could also be used to evaluate the effects of different reference genomes on alignment accuracy, such as comparing graph genomes containing different variant sets to each other and to linear references, as investigated using simulation in the FORGe study (Pritt *et al*., 2018).

The Rabema study of Holtgrewe et al. (2011) highlighted the disadvantages of evaluating aligners using only simulated reads and the correct-by-location definition. They developed the concept of the “trace tree” to enumerate mapping locations and a tool, Rabema, for computing all mappings of real or simulated sequencing reads less than a certain Hamming or edit distance. Aligners were evaluated on their ability to return all matches within the distance threshold, all best matches, or any best match. However, using a single truth set of the optimal locations of all matches leaves alignment heuristics and scoring functions as confounding factors. Also, fixing a maximum distance *k* disregards reads where the optimal match is further than *k* from the reference, which we show are the most error-prone in semiglobal alignment algorithms. Future benchmarking efforts comparing alignment algorithms, parameter settings, and graph versus linear reference genome paradigms can now be based on aligner-specific, real data computational gold standards generated using Vargas.

Alignment methods developers can use Vargas alignments to identify particular reads where the heuristic fails to find the optimal solution to the optimization problem posed, and can revise the heuristic strategies accordingly. Knowing optimal alignments for a subset of the input reads can serve as training data for identifying optimal alignment parameters, as tools like Teaser (Smolka *et al*., 2015) do using simulated reads. Such customization of parameters should be particularly effective for short (e.g. ChIP-seq) and/or error-prone (e.g. ancient DNA) reads.

Many short-read datasets use paired-end reads, where a DNA fragment is sequenced from both ends, typically with a few hundred bases between the read pairs. Heuristic aligners account for the fact that pairs should align concordantly to the reference, i.e. in a particular expected configuration based on the library preparation. Since concordance is not defined by the scoring function per se, and since checking for concordance of paired-end alignments can be implemented as a post-pass after each end has been aligned individually, we left paired-end alignment to future work. We evaluated Salmon’s RNA-seq alignments to the transcriptome, but Vargas’ optimization functions do not extend to spliced short-read alignment, such as aligning a RNA-seq read to a genome. Possible optimization functions for penalizing splicing events could depend on intron length, genomic nucleotides at the donor and acceptor sites, and a transcriptome annotation. Extending the dynamic programming model to find optimal solutions would require that every position in the read could be spliced to any pair of coordinates in the genome with a corresponding alignment penalty, exponentially increasing the possibilities to be explored. Vargas does not currently support the minimum-of-two-affine-functions gap scoring function used by Minimap2 (Li, 2018) or variation graphs that are not DAGs, which would limit its evaluation of an aligner that worked with variation graphs containing cycles, for example.

Because Vargas calculates all possible alignments of read to reference in the course of filling the dynamic programming matrix, every match above a certain minimum score could be reported. This could be useful in CRISPR guide RNA off-target analysis: current approaches are limited to a few mismatches; using Vargas would allow for a full edit distance scoring function and enumeration of distant alignments, including in the presence of genetic variation. Fully profiling all possible alignments of a read, or a substring extracted from the genome, could also be applied to the problem of characterizing mappability as explored in Lee and Schatz (2012) and Wilson et al. (2019).

## Acknowledgements

This work used the Extreme Science and Engineering Discovery Environment (XSEDE), which is supported by National Science Foundation grant number ACI-1548562. This research project was conducted using computational resources at the Maryland Advanced Research Computing Center (MARCC), and Texas Advanced Computing Center (TACC) at The University of Texas at Austin through allocation TG-DEB180021. We thank Daniel Baker for assistance in software development and Daehwan Kim for helpful comments on the manuscript.

## References

Aird, D. et al. (2011). Analyzing and minimizing PCR amplification bias in Illumina sequencing libraries. Genome Biology, 12(2), R18.

Ballouz, S. et al. (2019). Is it time to change the reference genome? Genome Biology, 20(1), 159.

Church, D. M. et al. (2015). Extending reference assembly models. Genome Biology, 16(1), 13.

Daily, J. (2016). Parasail: SIMD C library for global, semi-global, and local pairwise sequence alignments. BMC Bioinformatics, 17(1), 81.

Eberle, M. A. et al. (2017). A reference data set of 5.4 million phased human variants validated by genetic inheritance from sequencing a three-generation 17-member pedigree. Genome Research, 27(1), 157–164.

Farrar, M. (2007). Striped Smith-Waterman speeds database searches six times over other SIMD implementations. Bioinformatics, 23(2), 156–161.

Garrison, E. et al. (2018). Variation graph toolkit improves read mapping by representing genetic variation in the reference. Nature Biotechnology, 36(9), 875–879.

Gotoh, O. (1982). An improved algorithm for matching biological sequences. Journal of Molecular Biology, 162(3), 705–708.

Holtgrewe, M. et al. (2011). A novel and well-defined benchmarking method for second generation read mapping. BMC Bioinformatics, 12(1), 210.

Huang, L. et al. (2013). Short read alignment with populations of genomes. Bioinformatics, 29(13), i361–i370.

Intel Corporation (2015). Intel architecture instruction set extensions programming reference..

Jain, C. et al. (2019). Accelerating Sequence Alignment to Graphs. In 2019 IEEE International Parallel and Distributed Processing Symposium (IPDPS), pages 451–461. IEEE.

Jeffers, J. et al. (2016). Intel Xeon Phi Processor High Performance Programming: Knights Landing Edition. Morgan Kaufmann.

Kim, D. et al. (2019). Graph-based genome alignment and genotyping with HISAT2 and HISAT-genotype. Nature Biotechnology, 37(8), 907–915.

Langmead, B. (2017). A tandem simulation framework for predicting mapping quality. Genome Biology, 18(1), 152.

Langmead, B. and Salzberg, S. L. (2012). Fast gapped-read alignment with Bowtie 2. Nature Methods, 9(4), 357–359.

Lee, C. et al. (2002). Multiple sequence alignment using partial order graphs. Bioinformatics, 18(3), 452–464.

Lee, H. and Schatz, M. C. (2012). Genomic dark matter: the reliability of short read mapping illustrated by the genome mappability score. Bioinformatics, 28(16), 2097–2105.

Li, H. (2013). Aligning sequence reads, clone sequences and assembly contigs with BWA-MEM.

Li, H. (2018). Minimap2: pairwise alignment for nucleotide sequences. 34(May), 3094–3100.

Li, H. and Durbin, R. (2010). Fast and accurate long-read alignment with Burrows-Wheeler transform. Bioinformatics (Oxford, England), 26(5), 589–95.

Li, H. et al. (2008). Mapping short DNA sequencing reads and calling variants using mapping quality scores. Genome research, 18(11), 1851–8.

Li, H. et al. (2018). A synthetic-diploid benchmark for accurate variant-calling evaluation. Nature Methods, 15(8), 595–597.

Liu, Y. and Schmidt, B. (2014). SWAPHI: Smith-waterman protein database search on Xeon Phi coprocessors. In 2014 IEEE 25th International Conference on Application-Specific Systems, Architectures and Processors, pages 184–185. IEEE.

Liu, Y. et al. (2013). CUDASW++ 3.0: accelerating Smith-Waterman protein database search by coupling CPU and GPU SIMD instructions. BMC Bioinformatics, 14(1), 117.

Lowy-Gallego, E. et al. (2019). Variant calling on the GRCh38 assembly with the data from phase three of the 1000 Genomes Project. Wellcome Open Research, 4, 50.

Paten, B. et al. (2017). Genome graphs and the evolution of genome inference. Genome Research, 27(5), 665–676.

Patro, R. et al. (2017). Salmon provides fast and bias-aware quantification of transcript expression. Nature Methods, 14(4), 417–419.

Pritt, J. et al. (2018). FORGe: prioritizing variants for graph genomes. Genome Biology, 19(1), 220.

Rahn, R. et al. (2018). Generic accelerated sequence alignment in SeqAn using vectorization and multi-threading. Bioinformatics, 34(20), 3437–3445.

Rautiainen, M. et al. (2019). Bit-parallel sequence-to-graph alignment. Bioinformatics.

Rognes, T. (2011). Faster Smith-Waterman database searches with inter-sequence SIMD parallelisation. BMC bioinformatics, 12, 221.

Rognes, T. and Seeberg, E. (2000). Six-fold speed-up of Smith-Waterman sequence database searches using parallel processing on common microprocessors. Bioinformatics (Oxford, England), 16(8), 699–706.

Schneeberger, K. et al. (2009). Simultaneous alignment of short reads against multiple genomes. Genome Biology, 10(9), R98.

Smith, T. and Waterman, M. (1981). Identification of common molecular subsequences. Journal of Molecular Biology, 147(1), 195–197.

Smolka, M. et al. (2015). Teaser: Individualized benchmarking and optimization of read mapping results for NGS data. Genome Biology, 16(1).

Sodani, A. (2015). Knights landing (KNL): 2nd Generation Intel^®^ Xeon Phi processor. In 2015 IEEE Hot Chips 27 Symposium (HCS), pages 1–24. IEEE.

Tam, S. M. et al. (2018). SkyLake-SP: A 14nm 28-Core xeon^®^ processor. In 2018 IEEE International Solid - State Circuits Conference - (ISSCC), pages 34–36. IEEE.

Vijaya Satya, R. et al. (2012). A new strategy to reduce allelic bias in RNA-Seq readmapping. Nucleic Acids Research, 40(16), 1–9.

Wilson, L. O. W. et al. (2019). VARSCOT: variant-aware detection and scoring enables sensitive and personalized off-target detection for CRISPR-Cas9. BMC Biotechnol., 19(1), 40.

Wozniak, A. (1997). Using video-oriented instructions to speed up sequence comparison. Computer applications in the biosciences: CABIOS, 13(2), 145–50.

Yang, X. et al. (2019). One reference genome is not enough. Genome Biology, 20(1), 104.

Zook, J. M. et al. (2014). Integrating human sequence data sets provides a resource of benchmark SNP and indel genotype calls. Nature Biotechnology, 32(3), 246–251.

