## Supplementary Information for "Vargas: heuristic-free alignment for assessing linear and graph read aligners"

#### Datasets used

100bp WGS reads from the 1000 Genomes Project sample NA18505, SRA accession ERR239486 are available at `ftp://ftp.1000genomes.ebi.ac.uk/vol1/ftp/phase3/data/NA18505/sequence_read/ERR239486_1.filt.fastq.gz`. We use the first 100,000 reads.

250bp WGS reads from the 1000 Genomes Project sample NA19017, SRA accession SRR1295544. We used the first 100,000 reads accessed using sratoolkit fastq-dump (and analyzed Read 1 only since this is a paired-end library).

75bp RNA-seq reads from the GEUVADIS project, from 1000 Genomes Project cell line NA18505, SRA accession ERR204952 are available at `ftp://ftp.sra.ebi.ac.uk/vol1/fastq/ERR204/ERR204952/ERR204952_1.fastq.gz`. We use the first 100,000 reads.

41 million 36bp H3K27ac ChIP-seq reads from cell line GM18505, published in [10], SRA accession SRR901802 are available at `https://www.ncbi.nlm.nih.gov/sra/SRR901802`.

#### Simulation experiment (Related to Section 3.2)

Three sets of 100,000 unpaired reads of length 50, 100, and 150 bases respectively, were simulated with Mason2 [3] using the Illumina error model, human chromosome 19, with variants from the 1000 Genomes Project individual NA18505 [9]. Reads were aligned with Bowtie 2 and BWA-MEM to the chromosome 19 reference. The optimal alignment score for each aligner's scoring function was calculated by Vargas to compute correct-by-score. The simulated coordinate was compared to the aligned coordinate to calculate correct-by-location. Supplementary Table S4 lists the number of reads in each category.

Supplementary Table S4 indicates that very rarely, there are alignments that are incorrect-by-score but correct-by-location. For BWA-MEM, we determined that the optimal alignment computed by Vargas is shorter and applied soft-clipping to achieve a better score. For Bowtie 2, we determined that the optimal alignment computed by Vargas either has gaps near the end of the alignment which are disallowed by the Bowtie 2 `--gbar` parameter, or has mismatches at lower-base-quality bases than the mismatching bases in the simulated location, which led to a better score due to Bowtie 2's base quality scaled mismatch penalty.

#### Transcriptome alignment experiment (Related to Section 3.2)

The pseudoalignment and transcript quantification program Salmon [11] can optionally produce base pair level alignments with the `--writeMappings --validateMappings` command line flags. Since these alignments are generated with respect to an edit distance scoring function for semiglobal alignment and are reported with an alignment score, we can evaluate their accuracy with Vargas using the correct-by-score definition. The POS field corresponds to the pseudoalignment location, not the alignment score, so we can not use the correct-by-location definition. We used 100,000 RNA-seq reads described in the "Datasets" section above, and the Gencode version 33 transcriptome reference for GRCh38 for this experiment.

Vargas alignments of the reads to the transcriptome reference, (comprising 227,912 sequences totaling 359,080,835 bases) took 17.5 hours using 48 cores on a computer with AVX2 instructions, which allow for 32 reads per vector, but due to the scoring function and read length, 16-bit lanes were required which allowed for 16 reads per vector. We ran Salmon (custom version provided to us by the authors, which corrected a bug in alignment scoring present in release version 1.1.0) with and without the `--fullLengthAlignment` command line parameter which adjusts the heuristic strategy employed. The `--skipQuant` flag was also used which skips the quantification algorithm. Results are shown in Supplementary Table S5 and Supplementary Figure S6.

### Optimized alignment of ChIP-seq reads (Related to Section 4)

As an example of how Vargas alignments can be used to improve the heuristic alignment workflow for Bowtie 2, BWA-MEM and vg, we examined a set of 41 million unpaired 36-bp ChIP-seq reads from SRA accession SRR901802 [10]. In the study of [14], these reads were aligned with Bowtie 2 in an attempt to find novel QTLs. For initial alignment, we also used Bowtie 2 in the very-sensitive mode (runtime 6.5 minutes with 16 threads) and identified 570,000 (1.4%) reads that were unaligned or had an alignment with >1 mismatch or at least 1 gap. All further analysis was performed on this subset of the original dataset, which we refer to as difficult reads.

All of the difficult reads were realigned three times with Vargas, which was configured to match the default Bowtie 2 (semiglobal alignment), BWA-MEM, and vg scoring functions. vg alignment was performed to the graph of GRCh38 containing all 1000 Genomes Project variants with minor allele frequency > 10%; other alignments were performed to the linear GRCh38 reference. Alignment with Bowtie 2 parameters took 81 hours on 48 Skylake cores.

From these, we then selected 10,000 reads and aligned them with a selection of parameter settings for each of the three heuristic aligners. For Bowtie 2, we changed the seed length (-L) and dynamic programming extensions (-D) parameters; for BWA-MEM and vg we changed the seed length and reseeding parameters. Based on examining summary statistics comparing each parameter setting to the Vargas scores (Supplementary Excel File 2) we selected the combination of parameters for each tool that aligned at least as many reads as the default parameters and minimized the average difference in alignment score of all aligned reads. For BWA-MEM and Bowtie2, we also selected a parameter setting that took at most 1.5 times as long to run as the default setting. For vg, no parameter setting we explored met the additional runtime criteria. We realigned all the difficult reads with the new parameter settings. Results are summarized in Supplementary Table S6.

### Vargas memory consumption

The only sizable in-memory data structure used by Vargas is the reference genome. For the linear reference genome GRCh38, the representation consists of the string itself with one node per contig (199 total). On disk it is 2.9 Gb and alignment uses maximum 3.16 Gb. A graph genome requires a more complex representation, including the nodes and edges that represent various choices of alleles. The graph for GRCh38 plus all 1000 Genomes variants is 230 million nodes and 17 Gb on disk, with a total length linearized increase of 3.36% compared to the linear genome sequence. When the entire graph is loaded into memory, alignment uses maximum 101.3 Gb, although this could be reduced by changing the implementation to process the graph by chromosome or smaller chunks. The more practical graph used in the experiments in Section 3 and Section 4 with HISAT2 and vg contains the 1000 Genomes variants with MAF > 10%. It is 18.7 million nodes and 4 Gb on disk (0.24% length increase), and Vargas uses maximum 11 Gb memory in alignment.

### Trendline calculation

On all plots, the solid line shown for visualization purposes is fitted to the scatterplot using R's `geom_smooth` function with Loess smoothing and smoothing parameter 0.5. Note that the trendline is **not weighted** by the number of reads in each mapping quality or alignment score bin.

#### Vargas alignment parameters

**Bowtie 2 semiglobal and HISAT2** `--ete --ma 0 --mp 2,6 --np 1 --rdg 5,3 --rfg 5,3`

**Bowtie 2 local** `--ma 2 --mp 2,6 --np 1 --rdg 5,3 --rfg 5,3`

**BWA-MEM and vg** `--ma 1 --mp 4,4 --np 1 --rdg 6,1 --rfg 6,1`

**BWA aln** `--ete --ma 0 --mp 3,3 --np 3 --rdg 11,4 --rfg 11,4`

**Salmon** `--ete --ma 2 --mp 4,4 --rdg 4,2 --rfg 4,2 --np 0`

| | | | | All reads | | | Highest-scoring 95% | | | Lowest-scoring 5% | | |
|---|---|---|---|---|---|---|---|---|---|---|---|---|
| Tool | Mode | Time (s) | Mem. (GB) | % U | % I | % C | % U | % I | % C | % U | % I | % C |
| Bowtie 2 [6] semiglobal | very-sensitive | 100.89 | 3.38 | 1.03 | 0.63 | 98.35 | 0.01 | 0.15 | 99.84 | 20.59 | 12.09 | 67.32 |
| | sensitive (default) | 48.93 | 3.37 | 1.17 | 0.71 | 98.12 | 0.01 | 0.21 | 99.78 | 23.38 | 13.24 | 63.38 |
| | fast | 43.35 | 3.37 | 1.37 | 0.84 | 97.80 | 0.05 | 0.27 | 99.69 | 26.75 | 15.78 | 57.47 |
| | very-fast | 34.81 | 3.37 | 1.74 | 0.89 | 97.37 | 0.14 | 0.34 | 99.52 | 32.59 | 16.40 | 51.01 |
| Bowtie 2 [6] local | very-sensitive | 85.42 | 3.38 | 0.32 | 1.08 | 98.60 | 0.00 | 0.18 | 99.82 | 6.51 | 19.64 | 73.85 |
| | sensitive (default) | 64.28 | 3.38 | 0.33 | 1.26 | 98.41 | 0.00 | 0.26 | 99.74 | 6.71 | 21.84 | 71.44 |
| | fast | 38.32 | 3.37 | 0.38 | 1.52 | 98.11 | 0.00 | 0.32 | 99.68 | 7.65 | 26.51 | 65.84 |
| | very-fast | 32.74 | 3.37 | 0.49 | 1.85 | 97.66 | 0.01 | 0.48 | 99.52 | 9.84 | 31.29 | 58.87 |
| BWA-MEM [7] | k16 r1.2 | 87.51 | 5.51 | 0.35 | 0.43 | 99.23 | 0.00 | 0.04 | 99.96 | 7.05 | 8.52 | 84.43 |
| | k19 r1.5 (default) | 59.73 | 5.49 | 0.35 | 0.46 | 99.19 | 0.00 | 0.03 | 99.97 | 7.11 | 9.27 | 83.63 |
| | k22 r3 | 42.81 | 5.47 | 0.36 | 0.60 | 99.04 | 0.00 | 0.04 | 99.96 | 7.31 | 12.17 | 80.52 |
| | k25 r4 | 39.40 | 5.46 | 0.38 | 0.67 | 98.95 | 0.00 | 0.05 | 99.95 | 7.61 | 13.50 | 78.88 |
| BWA-MEM2 (AVX2) [15] | k16 r1.2 | 65.43 | 49.77 | 0.35 | 0.43 | 99.23 | 0.00 | 0.04 | 99.96 | 7.05 | 8.52 | 84.43 |
| | k19 r1.5 (default) | 75.29 | 49.66 | 0.35 | 0.46 | 99.19 | 0.00 | 0.03 | 99.97 | 7.11 | 9.27 | 83.63 |
| | k22 r3 | 69.51 | 49.62 | 0.36 | 0.60 | 99.04 | 0.00 | 0.04 | 99.96 | 7.31 | 12.17 | 80.52 |
| | k25 r4 | 67.78 | 49.60 | 0.38 | 0.67 | 98.95 | 0.00 | 0.05 | 99.95 | 7.61 | 13.50 | 78.88 |
| BWA aln [8] | o5 n15 | 245.67 | 4.25 | 1.38 | 0.42 | 98.21 | 0.01 | 0.03 | 99.97 | 28.24 | 11.11 | 60.65 |
| | o3 n10 | 253.87 | 3.87 | 1.63 | 0.33 | 98.03 | 0.01 | 0.02 | 99.97 | 33.48 | 9.50 | 57.01 |
| | o1 n5 (default) | 62.69 | 3.19 | 2.70 | 0.15 | 97.15 | 0.02 | 0.02 | 99.97 | 55.42 | 6.21 | 38.37 |
| HISAT2 [5] linear | very-sensitive | 44.09 | 4.47 | 0.64 | 1.38 | 97.98 | 0.69 | 1.48 | 97.83 | 12.16 | 24.82 | 63.01 |
| | sensitive | 25.43 | 4.47 | 1.56 | 0.96 | 97.48 | 1.56 | 1.04 | 97.40 | 29.66 | 17.55 | 52.79 |
| | fast (default) | 17.50 | 4.47 | 3.66 | 0.41 | 95.93 | 3.46 | 0.51 | 96.03 | 67.99 | 3.47 | 28.53 |
| HISAT2 [5] graph | very-sensitive | 52.73 | 6.73 | 0.69 | 1.37 | 97.95 | 0.08 | 0.26 | 99.66 | 12.58 | 22.81 | 64.60 |
| | sensitive | 58.06 | 6.73 | 1.56 | 0.93 | 97.51 | 0.12 | 0.29 | 99.59 | 29.64 | 13.40 | 56.96 |
| | fast (default) | 32.97 | 6.72 | 3.46 | 0.40 | 96.14 | 0.25 | 0.31 | 99.44 | 66.02 | 2.23 | 31.75 |
| vg [2] | linear | 397.20 | 24.16 | 0.19 | 0.56 | 99.25 | 0.00 | 0.04 | 99.96 | 3.81 | 10.59 | 85.60 |
| | graph | 381.63 | 26.58 | 0.19 | 0.51 | 99.31 | 0.00 | 0.03 | 99.97 | 4.24 | 10.89 | 84.87 |

Table S1: Alignment and correctness for the 100,000 100bp reads. Reported runtime is the median of three consecutive trials, and reported memory usage is the maximum memory footprint during alignment. U = unaligned; I = incorrect-by-score; C = correct-by-score. Time for bwa aln is reported for 'aln' only, not 'samse' which converts the intermediate output into SAM format. Related to Section 3.2.

| | | | | All reads | | | Highest-scoring 95% | | | Lowest-scoring 5% | | |
|---|---|---|---|---|---|---|---|---|---|---|---|---|
| Tool | Mode | Time (s) | Mem. (GB) | % U | % I | % C | % U | % I | % C | % U | % I | % C |
| Bowtie 2 [6] semiglobal | very-sensitive | 193.59 | 3.38 | 5.01 | 1.18 | 93.81 | 1.22 | 1.09 | 97.69 | 77.13 | 3.01 | 19.86 |
| | sensitive (default) | 97.23 | 3.37 | 6.11 | 1.39 | 92.50 | 2.25 | 1.29 | 96.46 | 79.56 | 3.47 | 16.97 |
| | fast | 69.15 | 3.37 | 7.42 | 1.89 | 90.70 | 3.50 | 1.79 | 94.71 | 81.92 | 3.73 | 14.35 |
| | very-fast | 47.98 | 3.37 | 8.38 | 2.45 | 89.17 | 4.40 | 2.40 | 93.20 | 84.17 | 3.49 | 12.35 |
| Bowtie 2 [6] local | very-sensitive | 237.80 | 3.40 | 2.14 | 2.26 | 95.60 | 0.54 | 1.15 | 98.31 | 32.93 | 23.71 | 43.35 |
| | sensitive (default) | 177.94 | 3.39 | 2.44 | 2.60 | 94.96 | 0.73 | 1.44 | 97.83 | 35.45 | 24.87 | 39.69 |
| | fast | 126.44 | 3.38 | 3.83 | 3.03 | 93.14 | 1.72 | 1.98 | 96.30 | 44.66 | 23.25 | 32.09 |
| | very-fast | 109.29 | 3.38 | 4.79 | 3.84 | 91.37 | 2.46 | 2.83 | 94.71 | 49.62 | 23.29 | 27.08 |
| BWA-MEM [7] | k16 r1.2 | 237.04 | 5.55 | 1.67 | 0.72 | 97.61 | 0.00 | 0.41 | 99.59 | 33.73 | 6.84 | 59.43 |
| | k19 r1.5 (default) | 119.74 | 5.45 | 1.84 | 0.86 | 97.30 | 0.01 | 0.46 | 99.53 | 36.92 | 8.64 | 54.44 |
| | k22 r3 | 97.70 | 5.43 | 2.27 | 1.11 | 96.63 | 0.60 | 0.60 | 98.80 | 43.88 | 10.82 | 45.30 |
| | k25 r4 | 89.74 | 5.43 | 2.78 | 1.23 | 95.99 | 0.26 | 0.70 | 99.04 | 51.12 | 11.32 | 37.56 |
| BWA-MEM2 (AVX2) [15] | k16 r1.2 | 187.52 | 50.04 | 1.67 | 0.72 | 97.61 | 0.00 | 0.41 | 99.59 | 33.73 | 6.84 | 59.43 |
| | k19 r1.5 (default) | 133.10 | 49.78 | 1.84 | 0.86 | 97.30 | 0.01 | 0.46 | 99.53 | 36.92 | 8.64 | 54.44 |
| | k22 r3 | 116.98 | 49.70 | 2.27 | 1.11 | 96.63 | 0.60 | 0.60 | 98.80 | 43.88 | 10.82 | 45.30 |
| | k25 r4 | 109.56 | 49.66 | 2.78 | 1.23 | 95.99 | 0.26 | 0.70 | 99.04 | 51.12 | 11.32 | 37.56 |
| HISAT2 [5] linear | very-sensitive | 124.43 | 4.48 | 3.46 | 3.53 | 93.01 | 1.36 | 2.46 | 96.18 | 43.33 | 23.99 | 32.69 |
| | sensitive | 61.76 | 4.48 | 6.95 | 2.82 | 90.23 | 2.37 | 2.92 | 94.71 | 94.05 | 0.88 | 5.07 |
| | fast (default) | 20.98 | 4.47 | 17.40 | 1.52 | 81.08 | 13.06 | 1.60 | 85.34 | 100.00 | 0.00 | 0.00 |
| HISAT2 [5] graph | very-sensitive | 171.59 | 6.74 | 3.47 | 4.09 | 92.43 | 1.37 | 3.00 | 95.63 | 43.58 | 24.88 | 31.54 |
| | sensitive | 96.25 | 6.73 | 6.96 | 3.24 | 89.80 | 2.40 | 3.37 | 94.23 | 94.23 | 0.87 | 5.00 |
| | fast (default) | 35.03 | 6.72 | 17.22 | 1.71 | 81.08 | 12.89 | 1.79 | 85.31 | 100.00 | 0.00 | 0.00 |
| vg [2] | linear | 633.29 | 24.15 | 1.31 | 1.29 | 97.41 | 0.06 | 0.44 | 99.50 | 25.34 | 17.47 | 57.19 |
| | graph | 692.07 | 26.55 | 1.29 | 1.30 | 97.41 | 0.06 | 0.44 | 99.50 | 24.78 | 17.68 | 57.54 |

Table S2: Alignment and correctness for the 100,000 250bp reads. Reported runtime is the median of three consecutive trials, and reported memory usage is the maximum memory footprint during alignment. U = unaligned; I = incorrect-by-score; C = correct-by-score. Related to Section 3.2.

| Tool | Reference | Availability | Type | Graph? |
|---|---|---|---|---|
| Vargas | this work | `https://github.com/langmead-lab/vargas` | Software | DAG |
| PaSGAL | [4] | `https://github.com/ParBLiSS/PaSGAL` | Software | DAG |
| vg align | [2] | `https://github.com/vgteam/vg` | Software | DAG |
| GraphAligner | [13] | `https://github.com/maickrau/GraphAligner` | Software | Any graph |
| SSW | [16] | `https://github.com/mengyao/Complete-Striped-Smith-Waterman-Library` | Library | no |
| GSSW | | `https://github.com/vgteam/gssw` | Library | DAG |
| seqan | [12] | `https://github.com/seqan/seqan3` | Library | no |
| Parasail | [1] | `https://github.com/jeffdaily/parasail` | Library | no |

| Tool | SIMD instructions | Vectorization strategy | Local? | Semiglobal? | Scoring function |
|---|---|---|---|---|---|
| Vargas | SSE, AVX2, AVX512BW | query-parallel | yes | yes | match/baseq-scaled mismatch<br>affine insertion/affine deletion |
| PaSGAL | AVX512BW | query-parallel | yes | no | match/mismatch/insertion/deletion |
| vg align | SSE | striped | yes | no | match/mismatch/affine gap |
| GraphAligner | no | bit-parallel | no | yes | unit cost |
| SSW | SSE | striped | yes | no | match/mismatch/affine gap |
| GSSW | SSE | striped | yes | no | match/mismatch/affine gap |
| seqan | SSE, AVX2, AVX512BW | query-parallel | yes | yes | match/mismatch/affine gap |
| Parasail | SSE, AVX2 | diagonal, blocked,<br>striped, prefix-scan | yes | no | match/mismatch/affine gap |

Table S3: Summary of exact dynamic programming pairwise alignment algorithms available in the literature and their features. Related to Section 3.1.

| Read length | Algorithm | Aligned reads | Correct-by-score + Correct-by-location | Incorrect-by-score + Correct-by-location | Correct-by-score + Incorrect-by-location | Inorrect-by-score + Incorrect-by-location |
|---:|:---:|---:|---:|---:|---:|---:|
| 50 | Bowtie2 | 99561 | 91502 | 34 | 7790 | 235 |
| 50 | BWA-MEM | 99947 | 91734 | 2 | 8080 | 131 |
| 100 | Bowtie2 | 99948 | 96961 | 23 | 2800 | 164 |
| 100 | BWA-MEM | 100000 | 97096 | 1 | 2871 | 32 |
| 150 | Bowtie2 | 99962 | 98513 | 17 | 1313 | 119 |
| 150 | BWA-MEM | 100000 | 98565 | 1 | 1431 | 3 |

Table S4: For simulated reads from human chromosome 19, comparison of correct-by-score and correct-by-location definitions. Optimal alignment score is calculated by Vargas; correct-by-location is calculated by comparing whether the simulated read location is equal to the aligned location. Related to Section 3.2.

| | | | | All reads | | | Highest-scoring 85% | | | Lowest-scoring 15% | | |
|---|---|---|---|---|---|---|---|---|---|---|---|---|
| Tool | Mode | Time (s) | Mem. (GB) | % U | % I | % C | % U | % I | % C | % U | % I | % C |
| Salmon [11] | default | 5.82 | 1.50 | 12.12 | 5.43 | 82.46 | 0.05 | 5.77 | 94.19 | 81.00 | 18.39 | 15.51 |
| | `--fullLengthAlignment` | 6.32 | 1.50 | 12.22 | 5.97 | 81.81 | 0.05 | 6.08 | 93.88 | 81.71 | 29.37 | 12.92 |

Table S5: Alignment and correctness for the 100,000 75bp RNA-seq reads aligned to the transcriptome. Reported runtime is the median of three consecutive trials with the –skipQuant flag, and reported memory usage is the maximum memory footprint during program execution. U = unaligned; I = incorrect-by-score; C = correct-by-score. Related to Section 3.2.

| | Bowtie2 SG | | BWA-MEM | | vg |
|---|---|---|---|---|---|
| Parameters | L14,D100 | L22,D100 | k11,r1.2 | k15,r1.5 | k10,r1.2 |
| % aligned | 1.27 | 1.01 | 1.06 | 1.05 | 1.25 |
| % correct score | 1.32 | 1.04 | 1.12 | 1.09 | 1.28 |
| Mean AS difference (all aligned reads) | 0.80 | 0.80 | 0.31 | 0.51 | 0.67 |
| Mean AS difference (incorrect-by-score aligned reads) | 0.99 | 0.93 | 0.76 | 0.87 | 1.12 |
| Alignment time / Default time | 14.63 | 1.76 | 8.14 | 2.33 | 14.59 |

Table S6: Bowtie 2 semiglobal alignment, BWA-MEM, and vg using MAF > 10% graph genome, were run on 570,000 difficult reads from the ChIP-seq dataset with various parameter settings. The value in the table denotes the ratio between the parameter setting shown and the the default parameters. For Bowtie 2 and BWA-MEM, the second parameter setting is the one that gave the lowest mean AS difference without aligning fewer reads or taking more than 1.5 times as long as the default setting in the 10,000-read experiment (Supplementary Excel File 2). Related to Section 4.

### References

[1] Daily, J. (2016). Parasail: SIMD C library for global, semi-global, and local pairwise sequence alignments. *BMC Bioinformatics*, **17**(1), 81.

[2] Garrison, E. *et al.* (2018). Variation graph toolkit improves read mapping by representing genetic variation in the reference. *Nature Biotechnology*, **36**(9), 875–879.

[3] Holtgrewe, M. (2010). Mason-A Read Simulator for Second Generation Sequencing Data. Technical report, Technical Report TR-B-10-06, Institut für Mathematik und Informatik, Freie Universität Berlin.

[4] Jain, C. *et al.* (2019). Accelerating Sequence Alignment to Graphs. In *2019 IEEE International Parallel and Distributed Processing Symposium (IPDPS)*, pages 451–461. IEEE.

[5] Kim, D. *et al.* (2019). Graph-based genome alignment and genotyping with HISAT2 and HISAT-genotype. *Nature Biotechnology*, **37**(8), 907–915.

[6] Langmead, B. and Salzberg, S. L. (2012). Fast gapped-read alignment with Bowtie 2. *Nature Methods*, **9**(4), 357–359.

[7] Li, H. (2013). Aligning sequence reads, clone sequences and assembly contigs with BWA-MEM.

[8] Li, H. *et al.* (2008). Mapping short DNA sequencing reads and calling variants using mapping quality scores. *Genome research*, **18**(11), 1851–8.

[9] Lowy-Gallego, E. *et al.* (2019). Variant calling on the GRCh38 assembly with the data from phase three of the 1000 Genomes Project. *Wellcome Open Research*, **4**, 50.

[10] McVicker, G. *et al.* (2013). Identification of Genetic Variants That Affect Histone Modifications in Human Cells. *Science*, **342**(6159), 747–749.

[11] Patro, R. *et al.* (2017). Salmon provides fast and bias-aware quantification of transcript expression. *Nature Methods*, **14**(4), 417–419.

[12] Rahn, R. *et al.* (2018). Generic accelerated sequence alignment in SeqAn using vectorization and multi-threading. *Bioinformatics*, **34**(20), 3437–3445.

[13] Rautiainen, M. *et al.* (2019). Bit-parallel sequence-to-graph alignment. *Bioinformatics*.

[14] Van De Geijn, B. *et al.* (2015). WASP: Allele-specific software for robust molecular quantitative trait locus discovery. *Nature Methods*, **12**(11), 1061–1063.

[15] Vasimuddin, M. *et al.* (2019). Efficient architecture-aware acceleration of BWA-MEM for multicore systems. In *Proceedings - 2019 IEEE 33rd International Parallel and Distributed Processing Symposium, IPDPS 2019*, pages 314–324. Institute of Electrical and Electronics Engineers Inc.

[16] Zhao, M. *et al.* (2013). SSW Library: An SIMD Smith-Waterman C/C++ Library for Use in Genomic Applications. *PLoS ONE*, **8**(12), e82138.

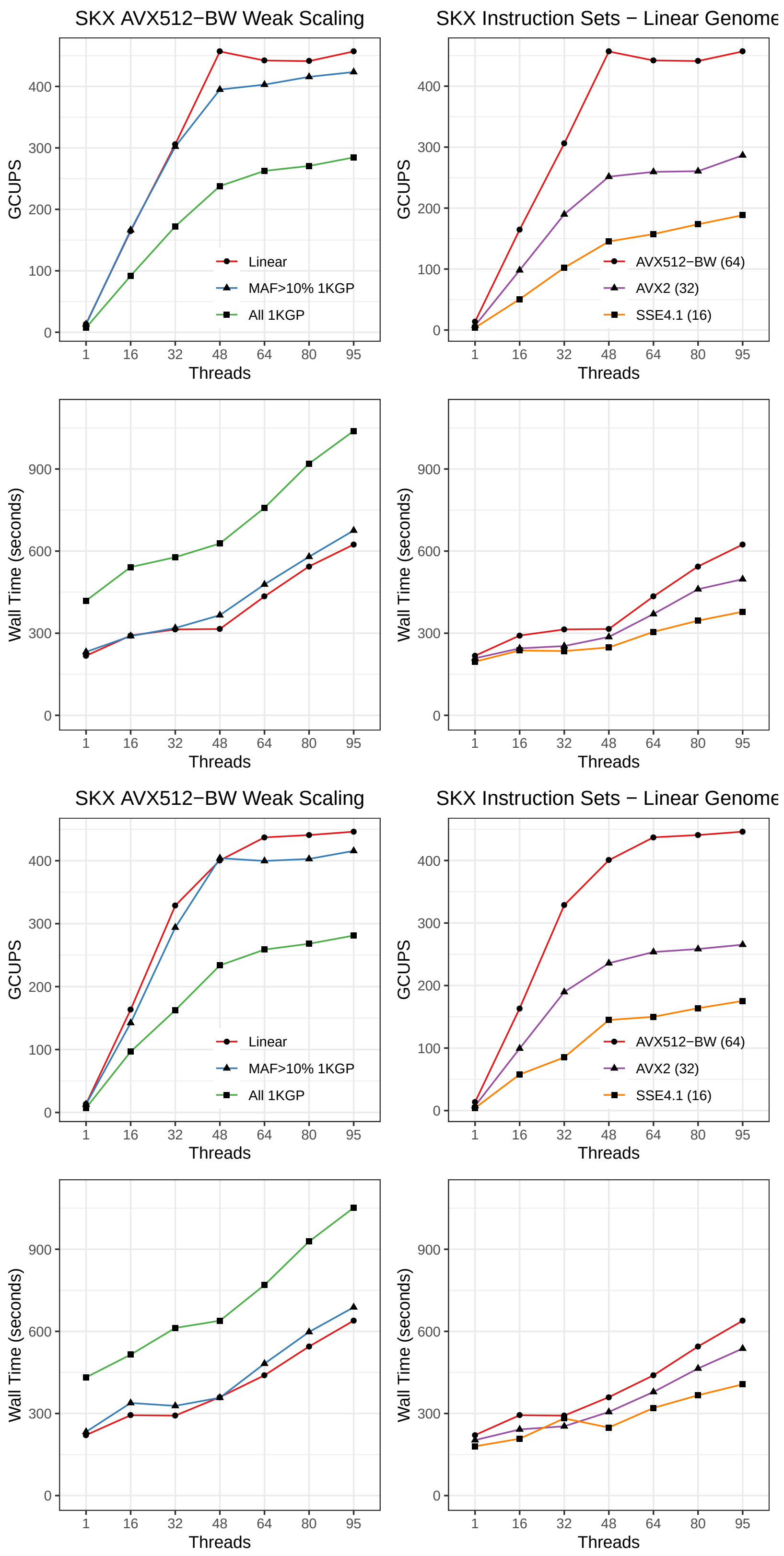

Figure S1: Weak scaling for Skylake (SKX), semiglobal alignment (top four) and local alignment (bottom four). Related to Figure 2.

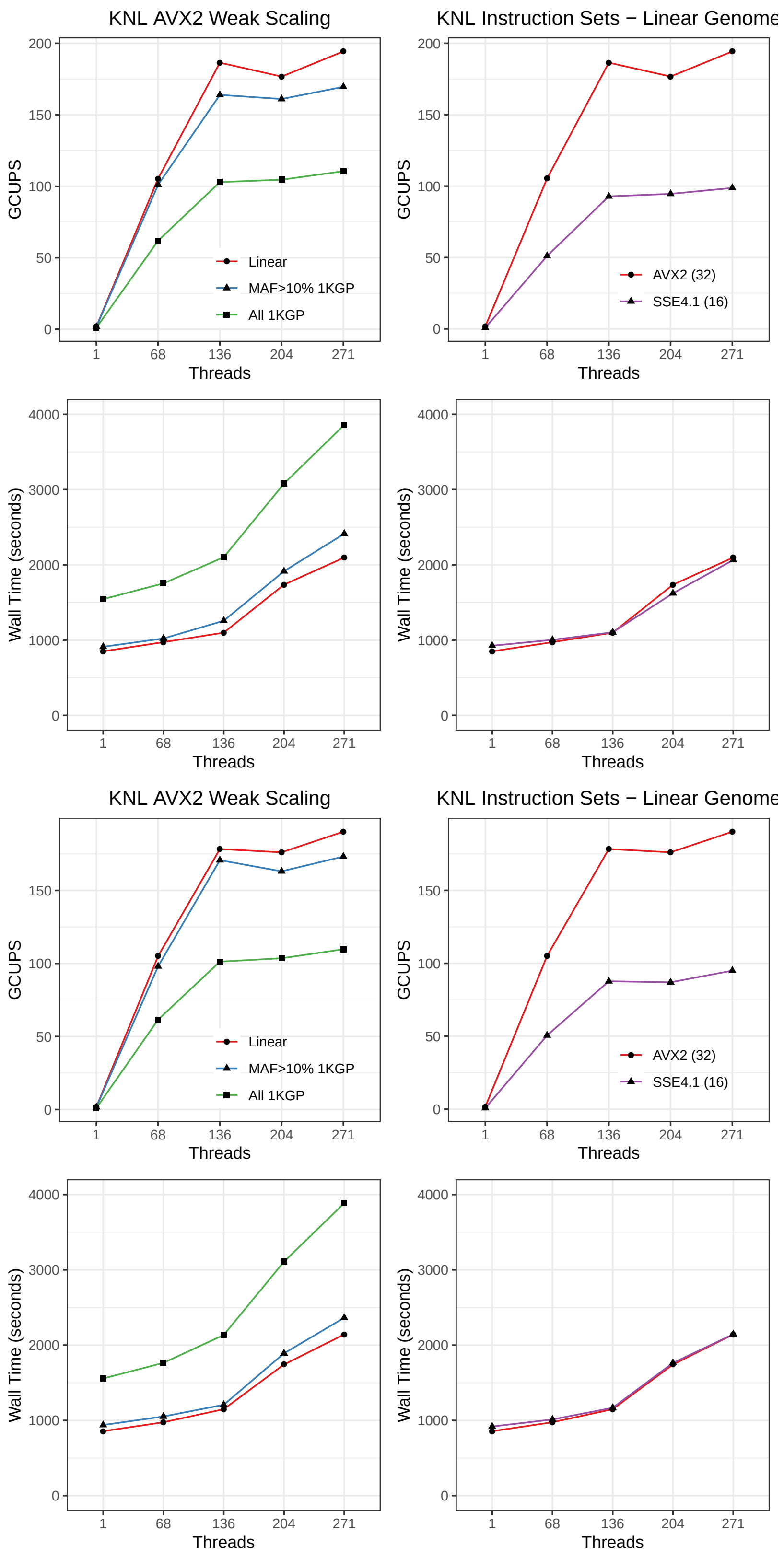

Figure S2: Weak scaling for Knight's Landing (KNL), semiglobal alignment (top four) and local alignment (bottom four). Related to Figure 2.

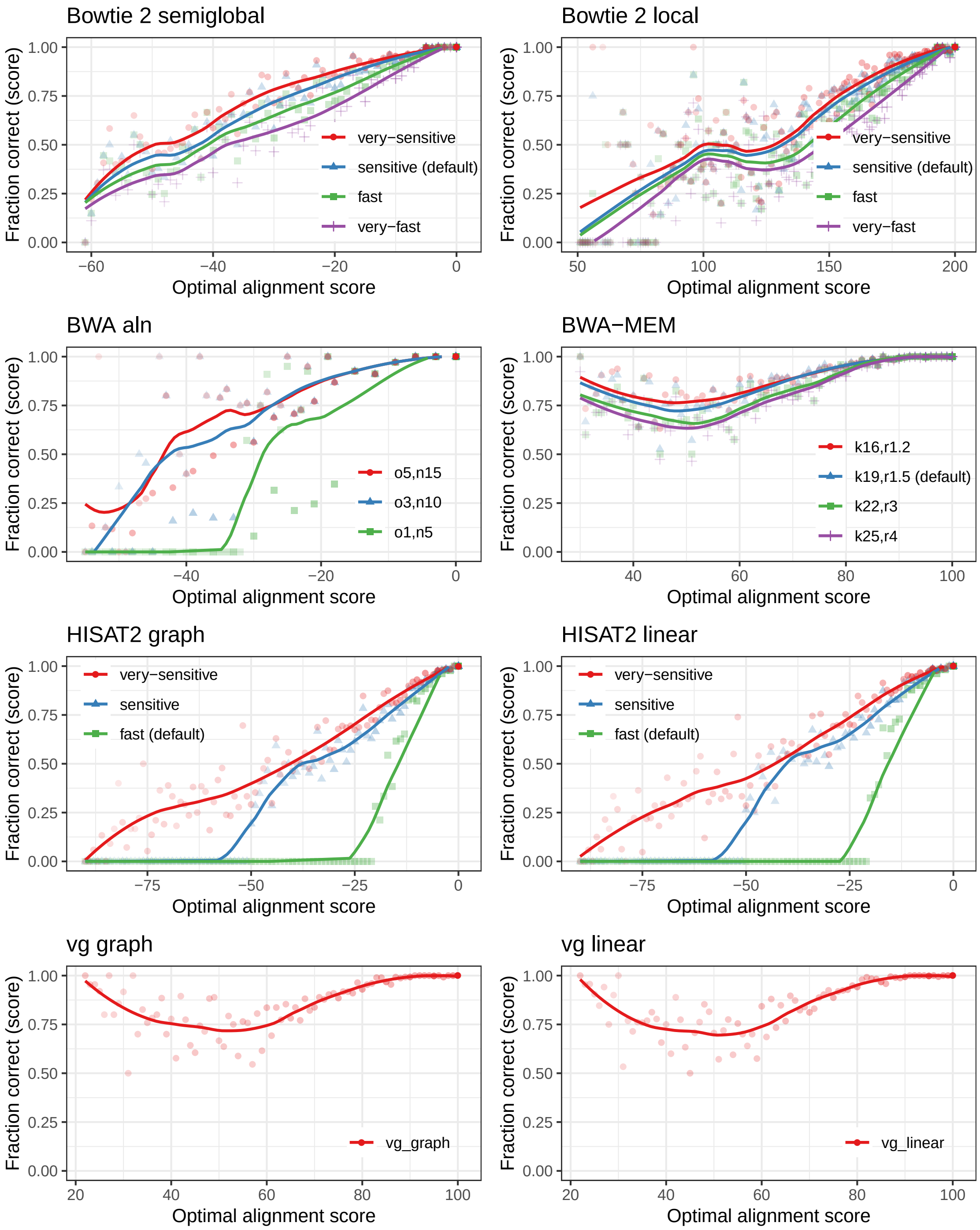

Figure S3: Correct-by-score plots for all aligners tested on the 100bp read dataset. Points are more transparent when representing fewer reads with a given optimal alignment score. Related to Figure 3.

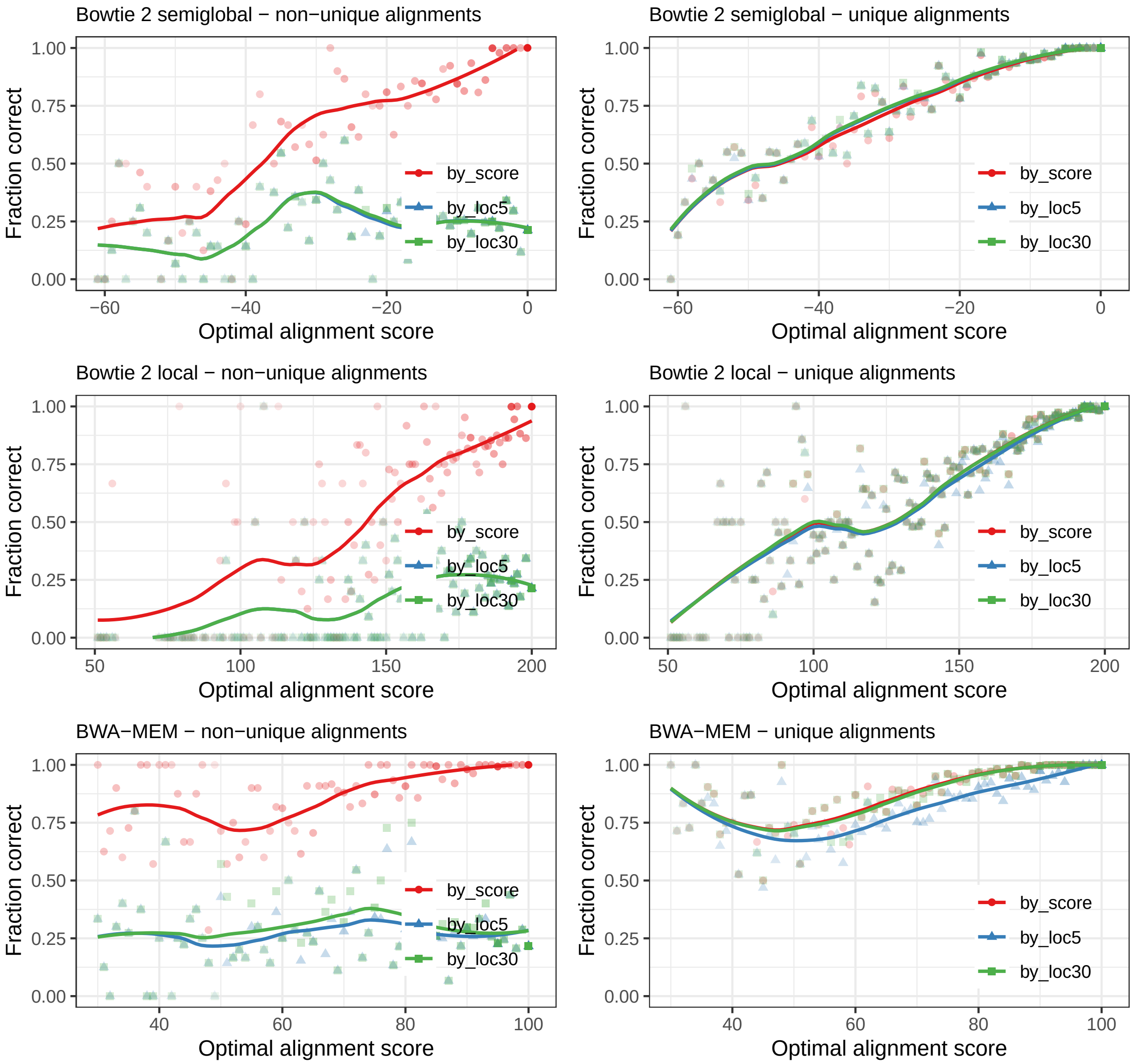

Figure S4: For the 100bp read dataset, comparing the correct-by-score and correct-by-location measurements, with different buffer sizes (0, 5, 30) for location of the left alignment coordinate. Since Vargas indicates whether the optimal alignment is unique within one read-length, the right column considers only reads that have a unique optimal alignment. Points are more transparent when representing fewer reads with a given optimal alignment score. Related to Figure 3.

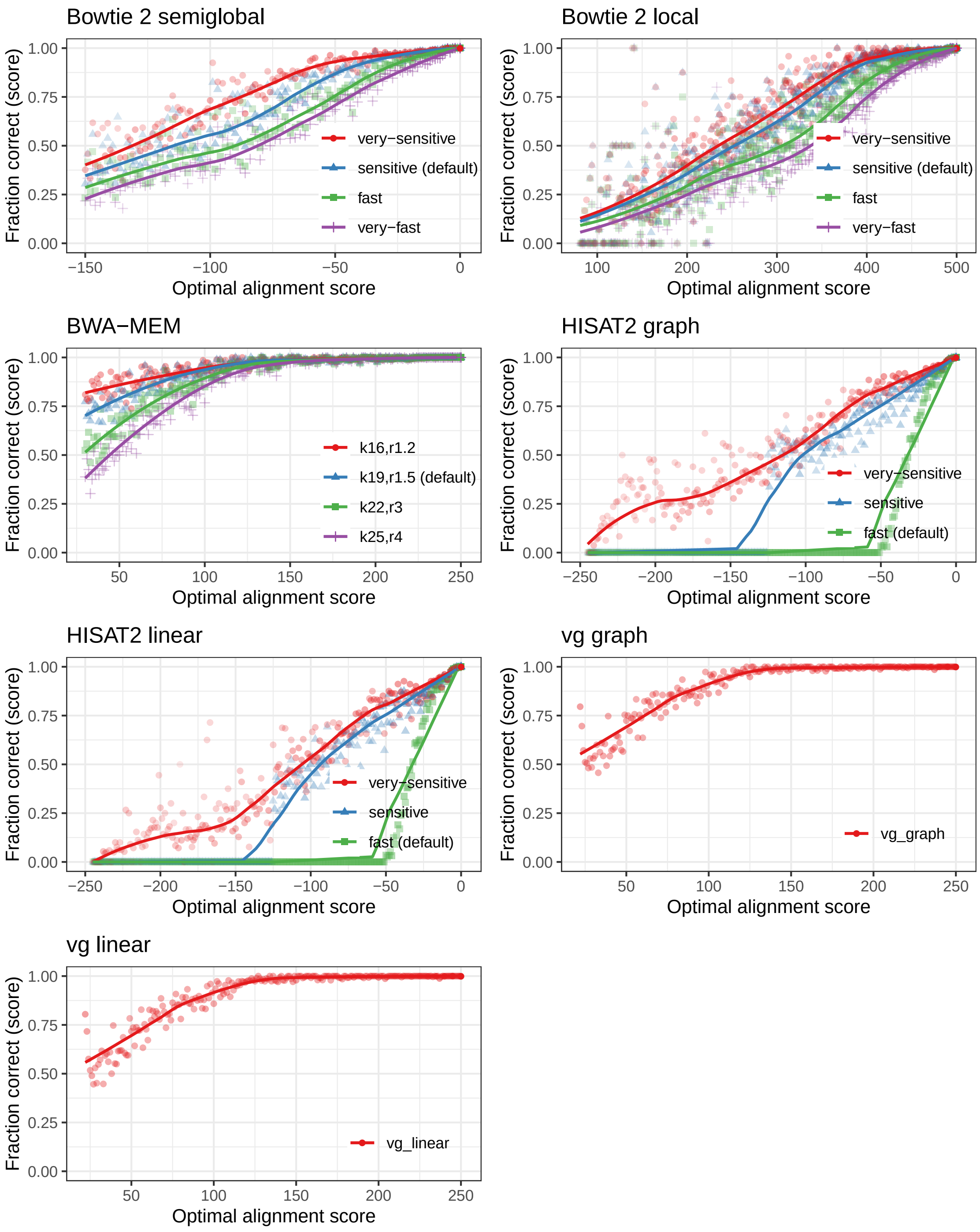

Figure S5: Correct-by-score plots for all aligners tested on the 250bp read dataset. Points are more transparent when representing fewer reads with a given optimal alignment score. Related to Figure 3 and Section 3.2.

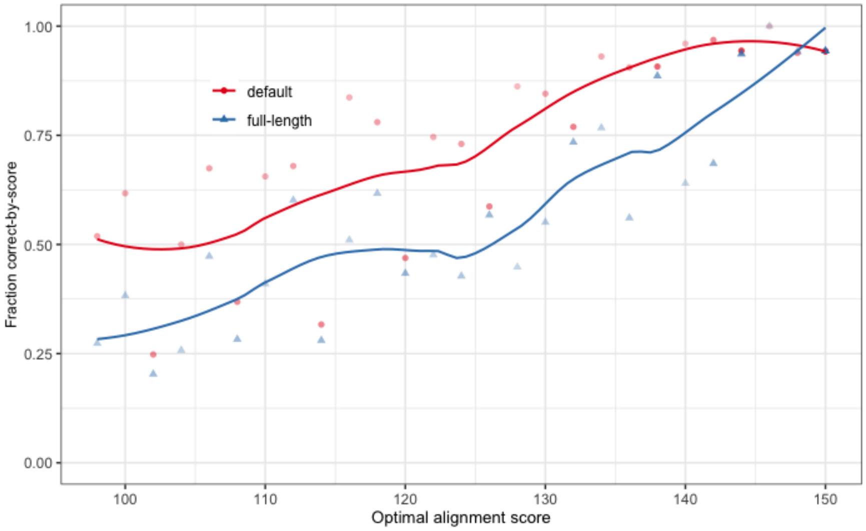

Figure S6: Correct-by-score plots for Salmon validation mappings. Points are more transparent when representing fewer reads with a given optimal alignment score. Related to Section 3.2.
